# Effects of Low-level Persistent Infection on maintenance of immunity by CD4 T cell subsets and Th1 cytokines

**DOI:** 10.1101/2022.12.01.518804

**Authors:** Samad A. Ibitokou, Komi Gbedande, Michael M. Opata, Victor H. Carpio, Karis M. Marshall, Robin Stephens

## Abstract

CD4 T cells are required, along with antibodies, for complete protection from blood-stage infection with *Plasmodium spp*., which cause malaria. Without continuous exposure, as on emigration of people from endemic areas, protection from malaria decays. As in other persistent infections, low-level *P. chabaudi* protects the host from re-infection at two months post-infection, a phenomenon termed premunition. Premunition is correlated with T cell responses, rather than antibody levels. We previously showed that while both effector T cells (Teff) and memory T cells (Tmem) are present after infection, Teff protect better than Tmem. Here we studied T cell kinetics post-infection by labelling dividing *Ifng^+^* T cells with BrdU in infected *Ifng-*reporter mice. A large drop in specific T cell numbers and *Ifng*^+^ cells upon clearance of parasite suggest a mechanism for decay of protection. Although protection decays, CD4 Tmem persist, including a highly-differentiated CD27^-^ Effector Memory (Tem) subset that maintains some *Ifng* expression. In addition, pre-treatment of chronically-infected animals with neutralizing antibody to IFN-γ, or clodronate liposomes before re-infection decrease premonition supporting a role for Th1-type immunity to re-infection. A pulse/chase experiment comparing chronically infected to treated animals showed that recently divided *Ifng^+^* T cells, particularly IFN-γ^+^TNF^+^IL-2^-^ T cells, are promoted by persistent infection. These data suggest that low-level persistent infection reduces CD4^+^ Tmem survival and multi-functional Teff but promotes IFN-γ^+^TNF^+^IL-2^-^ Late Effector Memory and Terminally Differentiated Effector T cells and prolongs immunity.

## Introduction

Malaria still accounts for 627,000 deaths globally, with an estimated 77% in children under 5 years of age (1). Morbidity from malaria is so prevalent that it has a measurable detrimental effect on the economic output of developing nations (2). *P. falciparum* can last up to one year in untreated people (3), and *P. chabaudi* infection lasts 2-3 months, a failure of this mostly protective response to eliminate parasite quickly. While sterile clearance is achieved in many acute infections, the late phase of low-level parasitemia does not induce fever or weight loss, suggesting co-evolutionary adaptation. Indeed, persistent *P. chabaudi* infection confers protection from re-infection, a phenomenon historically termed premunition, which is also seen in other persistent infections (4). Constant exposure protects people to some degree, as those who emigrate out of endemic areas lose protection, particularly from clinical disease, and become more ill than age-mates when infected again upon their return years later (5). This renewed susceptibility suggests that continuous exposure to the parasite promotes protection from malaria disease, and possibly parasitemia, however, adaptive mechanisms are unknown. Here, we studied costs and benefits of persistent infection to the adaptive immune system. *Plasmodium chabaudi* AS blood-stage infection has been well characterized (6). A maximum parasitemia occurs at days 8-10 post-infection (p.i.), which is largely resolved by day 20 p.i. followed by variable low-level recrudescence up to day 75 and sub-detectable levels lasting less than 90 days (7). Therefore, *P. chabaudi* infection of C57BL/6 mice represents a model of persistent *Plasmodium* infection that can be used to understand mechanisms of premunition (7).

Corresponding with the end of persistent infection, some immunity is lost, and understanding the cell types correlated with this protective phase and their decay can help us design better, longer-lasting vaccines. Malaria parasite-specific B cells and effector/memory-phenotype IFN-γ^+^ T cells accumulate with *P. falciparum* exposure, correlating with development of immunity (8, 9). *Plasmodium*-specific memory B cells have been shown to be stable in humans for up to 16 years, even without re-infection (10). However, the correlates of protection and survival of *Plasmodium*-specific memory CD4 T cells is only beginning to be understood (11, 12).

To determine correlates of protection in persistent infection, it is important to understand the kinetics of Th1 cell survival and effects of persistent infection on cytokine production in a physiologically relevant model. Our previous studies demonstrate that *P. chabaudi* infection produces a mixture of Teff and effector memory T cells (Tem) in the recrudescence phase (13). Despite the presence of memory T cells, transferred Teff protect from parasitemia more effectively. Our work suggests that MSP1-specific transgenic Teff activated by *P. chabaudi* infection protect immunodeficient mice from *P. chabaudi* infection much better than Tem (14). Clearance of persistent *P. chabaudi* infection at day 30 p.i. with anti-malarial drugs reduced the protection of premunition at day 60, but it did not change the ratio of central memory (Tcm) to Tem, suggesting that Tem are not responsible for the improved protection of premunition. However, we showed previously that removing live parasite in the premunition phase did reduce Th1 cytokines. IFN-γ^+^TNF^+^IL-2^-^ -producing CD62L^lo^ cells were correlated to prolonged protection, though we were unable to determine if they were Teff or Tem, which would be informative to understand how persistent infection maintains the protective Th1-type response (13). As chronic infection is protective, promotion of this particular cytokine producing population is suggestive that this T cell population promotes protection.

Understanding mechanisms underlying the durability of protection is also critical for children facing cyclic malaria seasons. Teff decay more quickly than Tem upon transfer into uninfected recipients, suggesting that they are short-lived. Although memory T cells develop, they appear to be unable to compete with parasite growth. Therefore, the decay of protective Teff is likely to explain the decay of immunity, but this has not been formally tested. Freitas do Rosario *et al*. showed in mice that it is the decay of *P. chabaudi*-responsive CD4 T cells, and not the level of parasite-specific antibody, that correlates with the decay of protection from parasitemia upon re-infection that occurs between days 120 and 200 p.i. (15). We recently showed that deletion of transcription factors in T cells to drive a more Th1 response promotes protection from re-infection (16). In addition, da Silva *et al.* show that recombinant IFN-γ can promote heterologous protection once T cell immunity decays at day 200 p.i. supporting a role for a Th1-type response in protection (17).

Here, we have studied the polyclonal response to *P. chabaudi* in order to measure their actual kinetics post-infection. We show that CD4 T cells are detectable in blood after *P. chabaudi* infection and decay in two waves. A dramatic decline of T cell numbers occurs after the first parasite peak, while Tmem numbers and a constant fraction of *Ifng*/*Thy1.1*^+^ in each subset are maintained more stably. Neutralization IFN-γ or depletion of phagocytic cells before reinfection reduced protection supporting a role for a Th1-type response in immunity. Studying T cell turnover, we identified Teff maintained by persistent infection that retained Ifng reporter expression, and showed that eliminating low level chronicity slightly reduced Ifng^+^ Teff. Recent proliferation was not particularly associated with *Ifng^+^* in Tmem and eliminating low level chronicity significantly increased survival of Tmem, but not the fraction of Tmem that were *Ifng*^+^. In addition, persistent infection reduced multi-functional IFN-γ^+^TNF^+^IL-2^+^ Teff, but promoted IFN-γ^+^TNF^+^IL-2^-^ T cells suggesting differential regulation of cytokine multi-producers by persistent stimulation in effector versus memory subsets. These data highlight the important role of Teff and the Th1 response in maintaining the protection from *Plasmodium* infection during low-level persistent parasitemia.

## Results

### CD4 effector T cells are detectable in blood after *P. chabaudi* infection and decay in two waves

*P. chabaudi* blood-stage infection of C57BL/6 mice is characterized by a large peak of parasitemia (days 8-10 p.i.) as shown in **Fig. 1A**,, which peaks before the acute symptomatic phase (days 10-14 p.i.), followed by persistent small recrudescences (detectable by blood smear up to day 30 p.i.) that are completely cleared between 2- and 3-months post-infection (7). Increased protection attributable to persistent infection, or premunition, is present through the first two months, and a further loss of protection can be measured by day 200 p.i. (15). The only correlate of premunition at day 60 so far is an increase in IFN-γ^+^TNF^+^IL-2^-^ CD62L^lo^ T cells in persistently infected mice compared to day 30 chloroquine-treated mice (12). It is not yet known if these are short lived effector T cells proliferating in response to persistent infection, or effector memory T cells.

**Figure 1.**
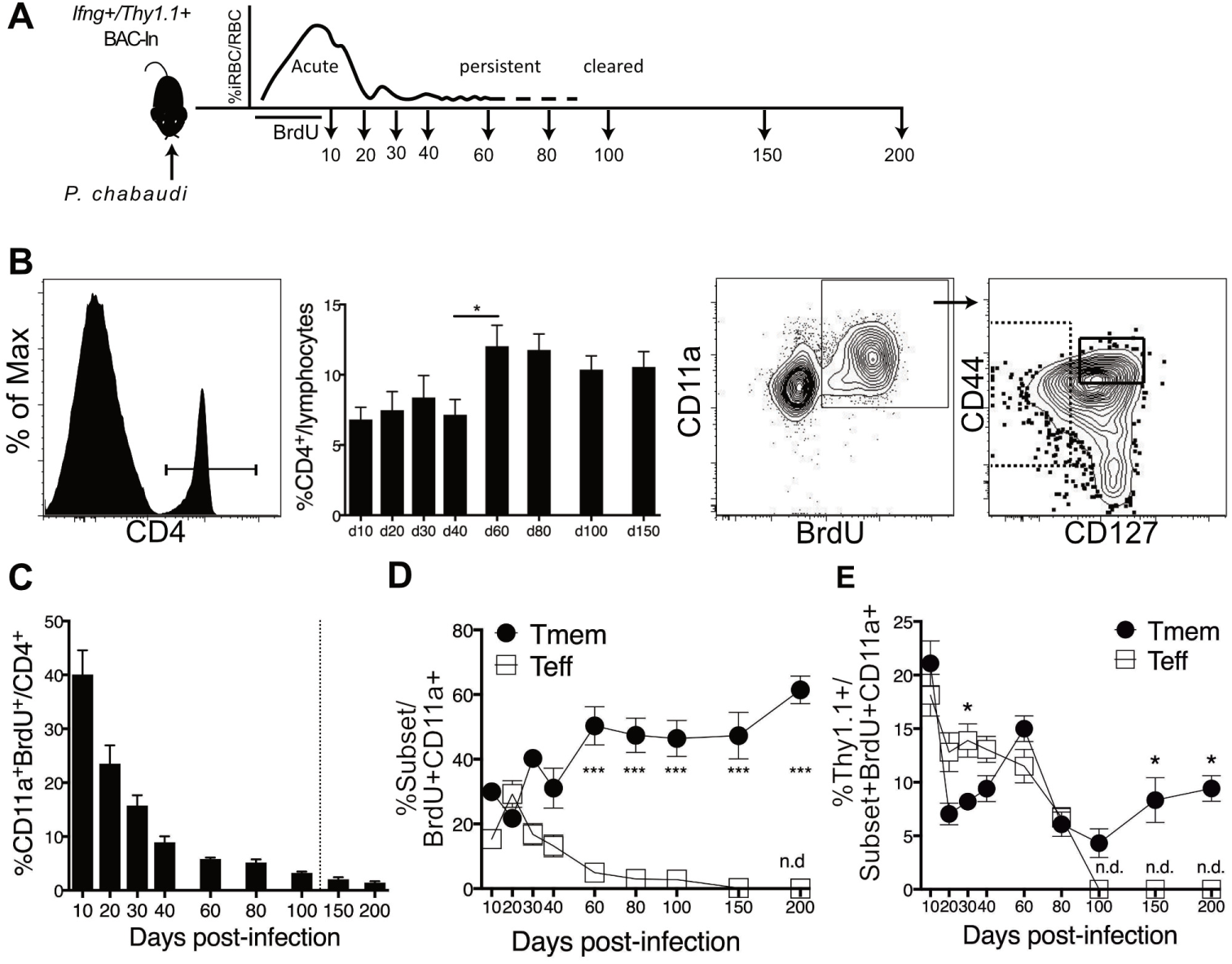
Decay of Teff and survival of Tmem after *P. chabaudi* infection. *Ifng*/Thy1.1 BAC-In reporter mice were infected with *P. chabaudi* and given BrdU from days 3-10 p.i. to label dividing cells. Blood was collected at indicated timepoints (days 10-200 p.i.) to study the decay of Teff and Tmem generated days 3-10 p.i. **(A)** Schematic representation of the experimental design. (**B**) Flow cytometry gating strategy showing percentage of CD4^+^ out of blood lymphocytes, CD4^+^CD11a^+^ BrdU^+^ Teff (CD127**^-^**) and Tmem (CD44^hi^CD127^+^). Graphs showing (**C**) percentage of CD11a^+^BrdU^+^ out of blood CD4 T cells, (**D**) fraction of Teff and Tmem within divided BrdU^+^CD11a^+^CD4^+^ T cells; and (**E**) the fraction within BrdU^+^CD11a^+^ Teff and Tmem that are of *Ifng*/Thy1.1^+^. Data are representative of three experiments with five mice per group. Error bars represent SEM, n.d., none detected, *p < 0.05, ***p < 0.001, Student’s *t* test.

To understand longevity and decay of CD4^+^ *Ifng*^+^ T cells during premunition, IFN-γ reporter mice were infected with *P. chabaudi* AS. Dividing *P. chabaudi*-responsive T cells in *Ifng*/Thy1.1 BAC-In mice were labelled with BrdU for seven days up to and including the peak of T cell expansion (**Fig. 1A**), on day 9 p.i. (18). CD4 T cells activated by infection were detected in the blood (**Fig. 1B**), by flow cytometry using antibodies to BrdU and CD11a, which has been shown to be upregulated in response to antigen, and not cytokines (19). As IL-7Rα (CD127) is completely downregulated upon activation of CD4 T cells, we used this as a marker to distinguish Teff from Tmem. Teff in the blood were defined as CD127^int/-^, in contrast to the spleen where the CD127^-^ population represents a discreet subset at the peak of infection (13). T cell activation was followed over 200 days using the fraction of CD4 T cells in the blood as a stable denominator to quantify the change of T cell subsets over time (**Fig. 1C**). Responsive T cells (CD4^+^BrdU^+^CD11a^+^) showed a dramatic decay, particularly between days 10 and 20 post-infection, and a slower consistent rate of decline after that. Specifically, only half of divided T cells remained by day 20 post-infection (p.i.) after T cell contraction, while some *Plasmodium*-specific T cells remained detectable in blood for 200 days. The fraction of CD127 ^int/-^ Teff of total CD11a^+^ BrdU^+^ CD4 in the blood was significantly decreased compared to that of memory T cells by day 60 p.i. (**Fig. 1D**). Teff, but not Tmem, became undetectable by day 200 p.i.., when protection from parasitemia has been shown to decay (15) (K. Gbedande, S. A. Ibitokou, M. L. Ong, M. A. Degli-Esposti, M. G. Brown, R. Stephens, Unpublished data) . The decay of *Ifng*^+^ T cells followed different kinetics with *Ifng*/Thy1.1*^+^* Teff still detectable until day 80 (**Fig. 1E**). BrdU^+^ Teff contain more *Ifng^+^* cells than Tmem on days 20-30 p.i, but BrdU^+^ *Ifng*^+^ Teff became undetectable in the blood by day 100. Some *Ifng*^+^ Tmem derived from the peak of infection (5-10%) remained *Ifng*^+^, even after parasite clearance.

### A fully differentiated Tem subset can maintain IFN-γ production

In our previous work, we have defined Teff and Tmem subsets of varying degrees of activation and differentiation respectively, using the markers CD27 and CD62L. CD62L^lo^ Teff are short-lived and make the most cytokines, while all Tmem subsets survive longer than the Teff (20). Teff^Late^ (CD127^-^ CD62L^lo^ CD27**^-^**) are the terminal subset, with a substantial fraction showing signs of early apoptosis or death (20). With these functional differences in mind, we evaluated the maintenance of the subsets of Teff and Tmem derived from the peak of infection, and their expression of *Ifng*. The proportion of Tmem-precursor Teff (Teff^Early^, CD62L^hi^ CD27^+^); and short-lived CD62L^lo^ Teff (SLEC), composed of intermediate Teff (Teff^Int^, CD27^+^) and Teff^Late^ (CD27^-^), out of the total BrdU^+^CD11a^+^CD4^+^ CD127^-^ Teff population was measured in the blood through 200 days p.i. (**Fig 2A**). All subsets decayed dramatically between days 10 and 40 p.i. Tmem (CD127^hi^ CD44^hi^) were also subdivided into central memory T cells (Tcm, CD62L^hi^ CD27^+^) and CD62L^lo^ Tem subsets (CD27^+^ Tem^Early^ and CD27^-^ Tem^Late^), which are listed here in the order of their level of differentiation (20). Tmem cells, surviving from the time of labelling during the peak of infection, maintained their proportions better than Teff over time, with cells in all Tmem populations remaining low but detectable on day 200 p.i. (**Fig 2B**). Therefore, polyclonal *Plasmodium*-responseive T cells with a memory phenotype do survive at physiological levels.

**Figure 2.**
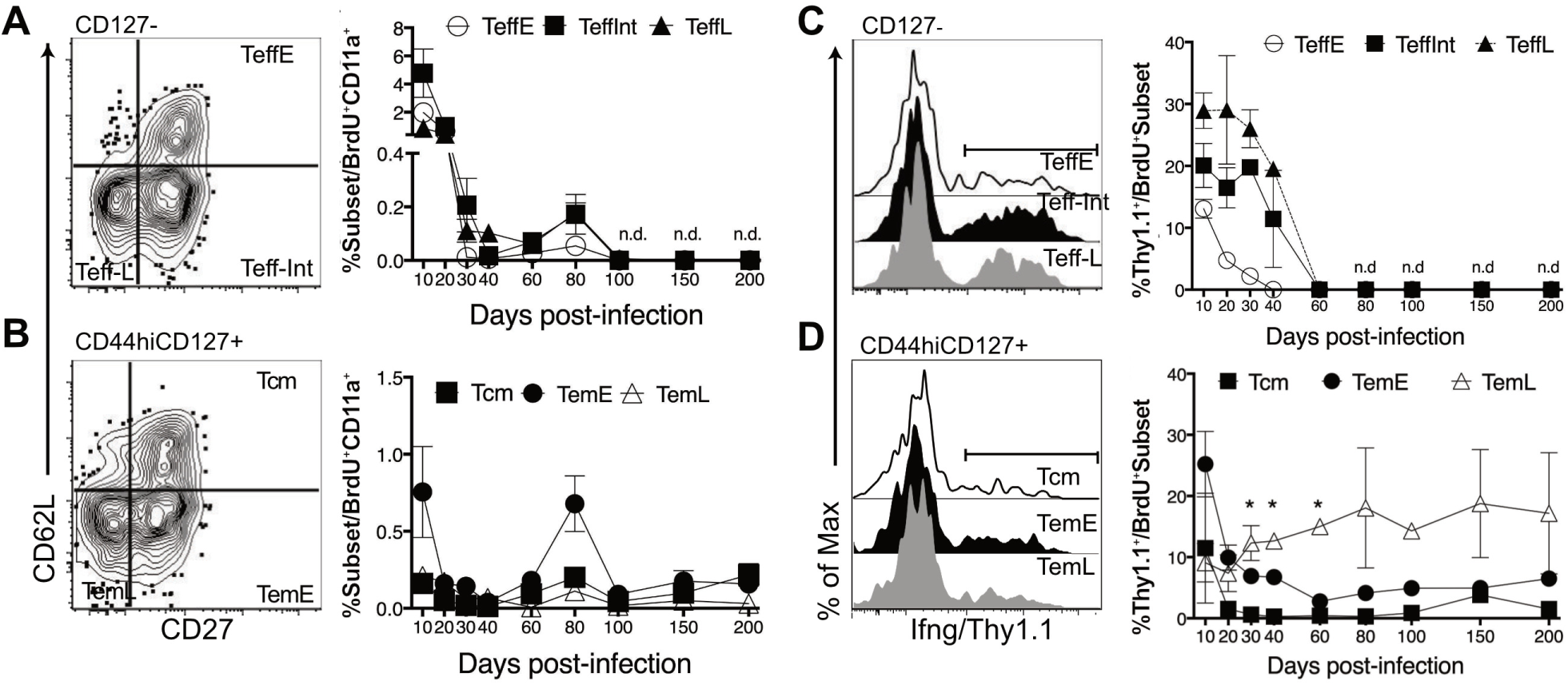
Decay of *Ifng* expression in *P. chabaudi* infection. *Ifng*/Thy1.1 BAC-In reporter mice were infected with *P. chabaudi* and given BrdU from days 3-10 p.i. to label dividing cells. Blood was collected (days 10-200) to study the decay cells labelled generated days 3-10 p.i including their current Teff and Tmem subset phenotypes (CD62L, CD27). (**A, B**) Concatenated contour plots with outliers and graphs showing the fraction of (**A**) each Teff (CD127^-^) subset (Teff^Early^: CD62L^hi^CD27^+^; Teff^Int^: CD62L^lo^CD27^+^ and Teff^Late^: CD62L^lo^CD27^-^) and (**B**) Tmem (CD44^hi^CD127^+^) subset (Tcm: CD62L^hi^CD27^+^; Tem^Early^: CD62L^lo^CD27^+^ and Tem^Late^: CD62L^lo^CD27^-^) that are BrdU^+^ at each timepoint. (**C, D**) Histograms showing percentage of CD11a^+^BrdU^+^ (**C**) Teff and (**D**) Tmem subsets expressing *Ifng*/Thy1.1^+^. Error bars represent SEM, n.d., none detected, *p < 0.05 *Ifng*^+^ TemL compared to Tcm and TemE.

IFN-γ production by Th1 cells, and *Ifng* BAC-In reporter expression, can be stable after acute infection (21–26), but this is less well established in persistent infection where exhaustion is common (27). In order to test for maintenance of *Ifng*, we observed expression of the *Ifng*/Thy1.1 Bac-In reporter on BrdU^+^ T cells in the blood through day 200. Despite this reporter including an RNA-stabilizing SV40 poly A tail, *Ifng/*Thy1.1 reporter became undetectable in all Teff subsets in the blood at day 80 p.i., significantly earlier than the Teff numbers decreased below detection (**Fig 2C**). Mature Teff (CD62L^lo^ Teff^Int^ and Teff^Late^) maintained a high, and relatively constant, fraction (18-30%) of *Ifng*/Thy1.1^+^ T cells from 10 to 30 days p.i., while Teff^Early^ showed a transient peak of *Ifng* at day 10. In Tmem subsets, *Ifng*/Thy1.1^+^ cells were detected up to day 200 p.i. (**Fig 2D**). The most differentiated subset of memory T cells, CD27-Tem^Late^, maintained the highest fraction of *Ifng*/Thy1.1^+^, significantly more than Tcm and Tem^Early^ at three timepoints. Together, these data suggest that the fraction of *Ifng^+^* in fully activated Teff subsets is quite constant until day 40, while highly differentiated Tem^Late^ maintain constant *Ifng* expression through day 200. Notably, the disappearance of *Ifng^+^* Teff in blood by day 80 corresponds to the complete clearance of the parasite, which we have documented to occur between days 75-90 p.i. (7).

### Teff numbers and *Ifng* expression are promoted during premunition in the presence of parasite

Loss of detectable *Ifng*^+^ Teff cells from the blood could represent their death, their migration into tissues, or their transition into Tmem. In this blood-borne infection, the tissue affected is the spleen. Therefore, to probe migration and generation of Tmem from Teff cells that proliferated during the peak of infection, we quantified BrdU^+^ *Ifng^+^* Teff and Tmem in the spleen. Infected animals that received BrdU on days 3-10 p.i. had splenocytes harvested, and the proportion of BrdU^+^ out of CD4^+^ and their and number were measured on days 60, where premunition is detectable, and 120 p.i., where it is not, up to day 200 p.i., when protection from parasitemia has been shown to be detectably lost (15). A decline of over half of the fraction and number of previously proliferated CD4 T cells occurred between days 60 and 120 (**Fig 3A**). The decrease in total BrdU^+^ T cells from day 60 to day 120 is accounted for by the decrease in BrdU^+^ Teff numbers, not memory (**Fig 3B**). There were significant proportions and numbers of BrdU^+^ Tmem (CD127^hi^ CD44^hi^) and Teff (CD127^-^) above the threshold of detection in the spleen through day 200, but more Tmem at all of these late timepoints. Gating within *Ifng*/Thy1.1^+^ shows that the number of *Ifng^+^* Tmem in the spleen was also significantly higher than *Ifng^+^* Teff on days 120 and 200 p.i., when *Ifng^+^* Teff disappear below the limit of detection in the spleen (**Fig 3C**). These data indicate that very few *Ifng*-expressing Teff derived from the peak of infection are maintained through day 120 after infection with *P. chabaudi*. Therefore, we hypothesized that T cells proliferating in response to the low-level persistent phase before day 60, contribute to the *Ifng*^+^ effector T cell population and to premunition which fall off after day 60.

**Figure 3.**
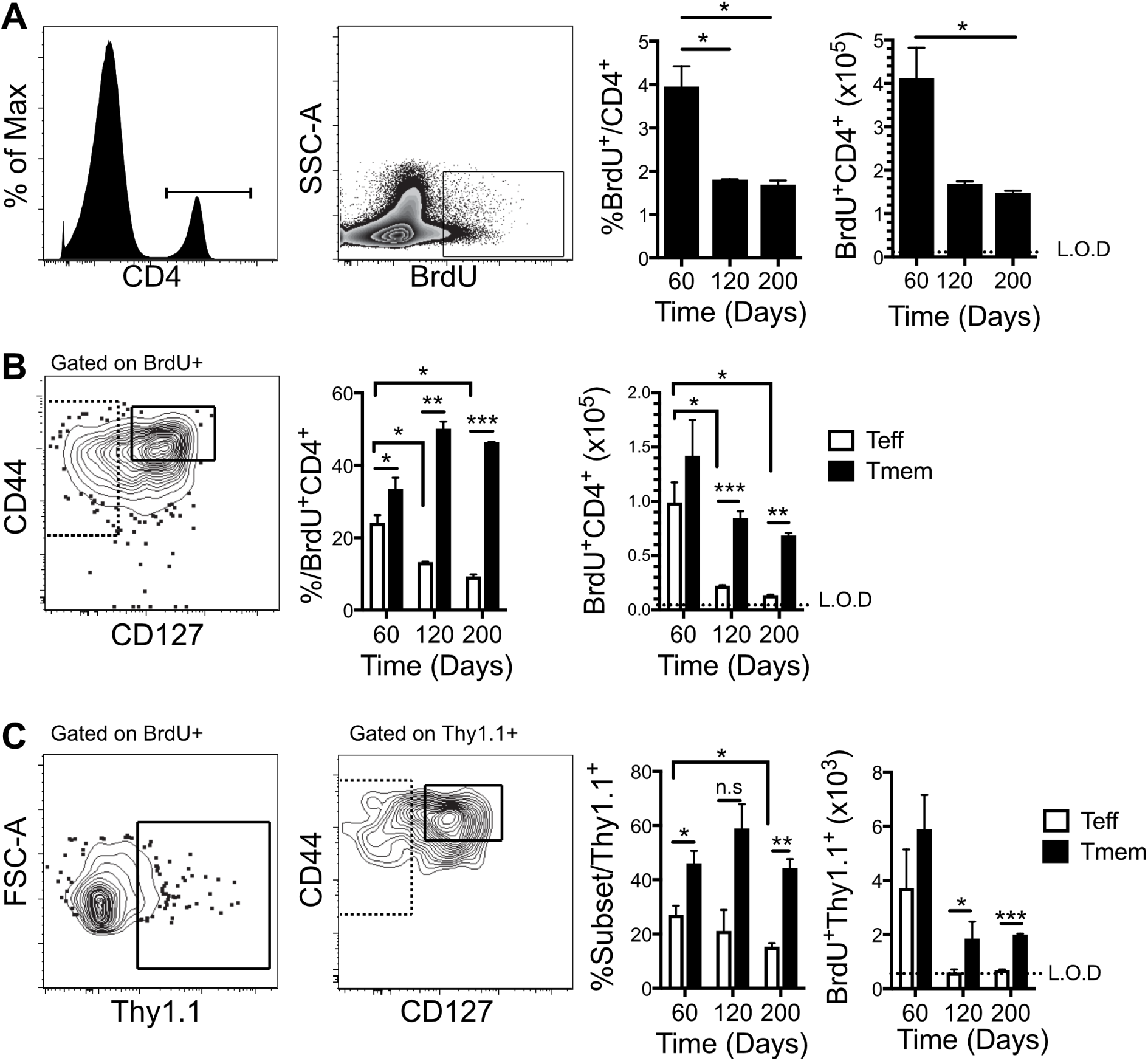
Decay of *Ifng*^+^ Teff between days 60 and 200 p.i. corresponds with decay of protection from heterologous infection. *Ifng*/Thy1.1 BAC-In reporter mice were infected with *P. chabaudi* and given BrdU from days 3-10 p.i. to label dividing cells and splenocytes analyzed (days 60, 120 and 200 p.i.). (**A)** Histogram and zebra plot show gating strategy, and graphs showing percentage of splenic CD4 T cells that are still BrdU^+^, and number of BrdU^+^ CD4^+^ T cells recovered at each timepoint. (**B**) Concatenated contour plot of BrdU+ T cells shows gating strategy, and graphs showing the fraction of Teff (CD127**^-^**) and Tmem (CD44^hi^CD127^hi^) out of BrdU^+^CD4^+^, and the number of BrdU^+^ Teff and Tmem. (**C**) Contour plots show gating strategies for determining *Ifng*/Thy1.1^+^ out of BrdU^+^, and graphs showing the fraction of BrdU^+^ *Ifng*/Thy1.1^+^ surviving as Teff and Tmem or total number of BrdU^+^ *Ifng*/Thy1.1^+^ Teff and Tmem. The limit of detection (L.O.D) was calculated using all available data from uninfected animals from all experiments that did not receive BrdU. Data are representative of three experiments with five mice per group. Error bars represent SEM, *p < 0.05, **p < 0.01, ***p < 0.001, Student’s *t* test.

### Teff and Tmem differentiation and survival in chronic *P. chabaudi* infection

To investigate the possibility that continuous generation of effector T cells contributes to protection during premunition, we labelled T cells with BrdU *in vivo* at later time points, for ten days starting either at day 20 or day 50 p.i., as shown in **Figure 4A**, and studied the survival of T cells that had divided (BrdU^+^) before (d20-30 p.i.) and after (d50-60) parasitemia becomes undetectable by slide counting (L.O.D. <%.001 iRBC/RBC). T cells were measured on day 60 p.i., when premunition is detectable. Interestingly, equal numbers of CD4^+^ BrdU^+^ (**Fig 4B**) and *Ifng*/Thy1.1^+^ BrdU^+^ (**Fig 4C**) T cells were observed at day 60, whether labelled on days 20-30, or 50-60 p.i. There was a trend toward more *Ifng*^+^BrdU^+^ T cells in the more recently labelled group (d50-60) compared to those labelled a month prior; however, this did not reach significance. There was also a slight but non-significant increase in Teff cells and *Ifng*^+^ Teff, but not Tmem, in more recently divided T cells (d50-60, **Fig 4D**). Within the BrdU^+^ effector and memory T cells that were generated either in the ten days before day 30 or day 60 p.i., and still present at day 60 p.i., there were mostly CD62L^lo^ cells (**Fig 4E**). There was a distinct and significant increase in *Ifng*/Thy1.1^+^ among mature Teff subsets that were more recently labeled (d50-60, **Fig 4F**). These data support the conclusion that *Ifng* expression is maintained in mature Teff by recent stimulation, as first noted in adoptive transfer of reporter cells (18). In addition, the data suggest quite a short timeframe for maintenance of *Ifng* accessibility in the absence of persistent infection, despite poly A tail driven stabilization of the BAC-In *Ifng*/Thy1.1 mRNA (22).

**Figure 4.**
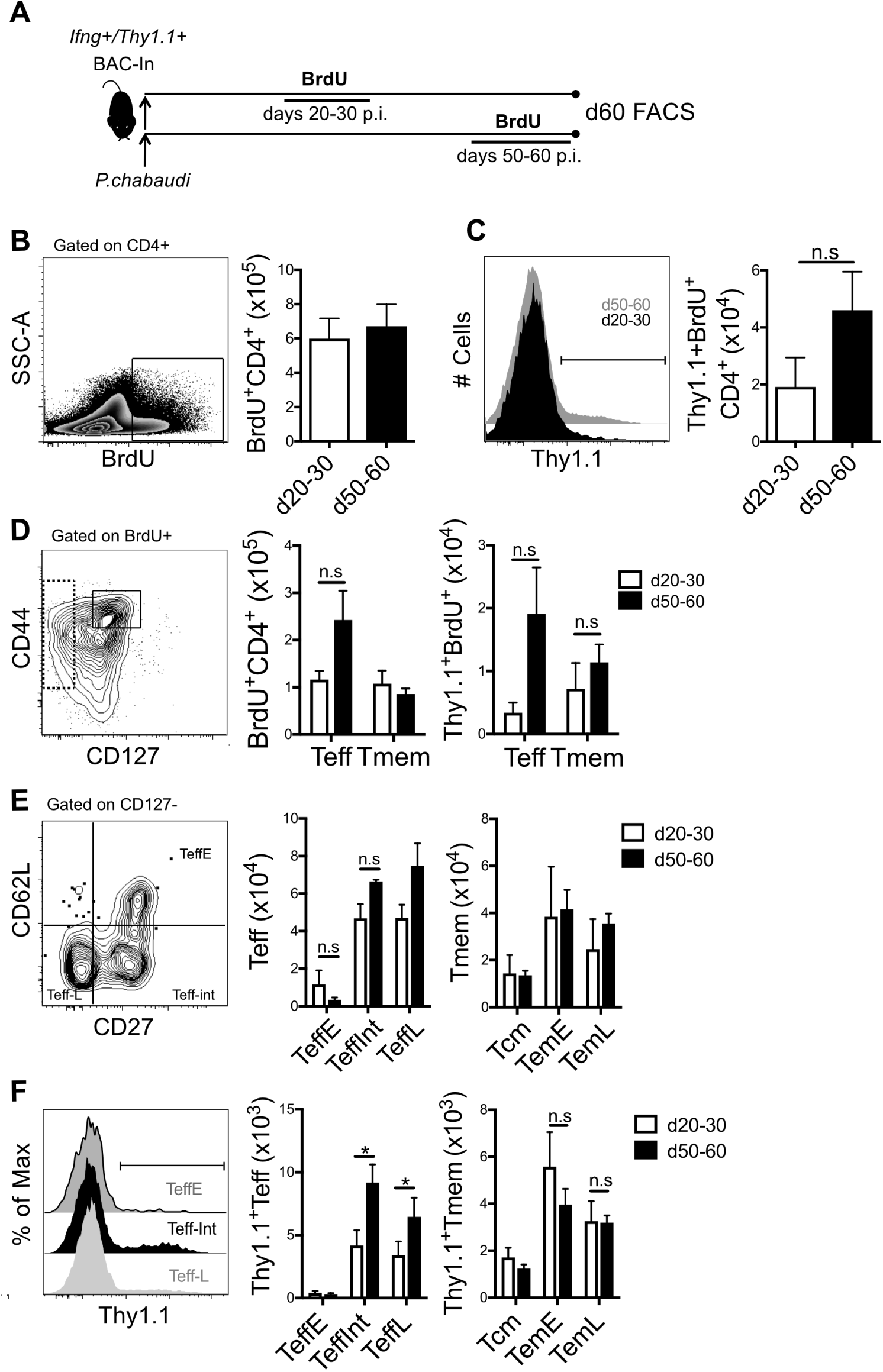
*Ifng*^+^ Teff are promoted by recent infection. (**A**) Schematic showing that *Ifng*/Thy1.1 BAC-In reporter mice were infected with *P. chabaudi* and given BrdU from days 20-30 or 50-60 p.i. Splenocytes were harvested at day 60 p.i. in both groups to study the day 60 phenotype of T cells generated during days 20-30 or 50-60, and their decay of *Ifng* expression. **(B)** Contour plots of BrdU^+^ out of CD4^+^ and graph of number of BrdU^+^CD4^+^ generated within each timeframe surviving until d60. (**C**) Histogram showing representative example of *Ifng*/Thy1.1^+^ within BrdU^+^CD4^+^ and graph the number of *Ifng*/Thy1.1^+^ BrdU^+^CD4^+^ generated with each timeframe observed at day 60. (**D**) Gating strategy showing representative example of Tmem (CD44^hi^CD127^hi^) Teff (CD127-) subsets from BrdU^+^CD4^+^ T cells and graphs showing numbers of BrdU^+^ Teff and Tmem and *Ifng*/Thy1.1^+^ Teff and Tmem generated with each timeframe observed at day 60. (**E**) Gating strategy showing representative example of Teff subsets to define Early Teff (Teff^Early^, CD62L^hi^ CD27^+^) and intermediate Teff (Teff^Int^, CD62L^lo^ CD27^+^) and late Teff (Teff^Late^, CD62L^lo^ CD27^-^) and Memory subsets (not shown): central memory T cells (Tcm, CD62L^hi^ CD27^+^) and effector memory T cells (CD62L^lo^CD27^+^ Tem^Early^ and CD62L^lo^CD27^-^Tem^Late^). Graphs show numbers of cells in each of these subsets that were generated in each timeframe of BrdU labelling and survive. (**F**) Histograms showing representative example of *Ifng*/Thy1.1^+^ within subsets and graphs showing the numbers of *Ifng*/Thy1.1^+^ cells generated at days 20-30 or 50-60 with Teff and Tmem subset phenotypes on day 60. Error bars represent SEM, n.s., not significant, *p < 0.05, Student’s *t* test.

### IFN-γ in premunition and Immune cell types mediating protection

As expression and decay of Ifng correlated with persistent infection, we tested the hypothesis that promotion of IFN-γ production by persistent infection is a mechanism for enhanced parasite immunity during chronic infection, and investigated the relative importance of different potential IFN-γ secreting cells on protection from re-infection during premunition. C57BL/6J mice were infected with *P. chabaudi* AS and premunition to heterologous re-infection was tested on day 60 p.i.. Some infected animals (n=5/group) were administered anti-IFN-γ neutralizing antibody or recombinant IFN-γ (rIFN-γ) or isotype over an eight-day period (d52-60 p.i.) before heterologous challenge with *P. chabaudi* AJ at day 60 post-primary infection (**Figure 5A**). Animals persistently infected with the AS clone were able to control heterologous challenge by day 6 suggesting premunition has an effect on heterologous infection (**Fig 5B**). We did not detect improvement of the excellent clearance observed in chronically-infected mice upon pre-treatment with rIFN-γ. However, if IFN-γ was blocked prior to re-infection during chronic infection, parasitemia was still uncontrolled, near 1.0%, after 8 days. Therefore IFN-γ is essential to the protective effect induced by chronic infection known as premunition.

**Figure 5.**
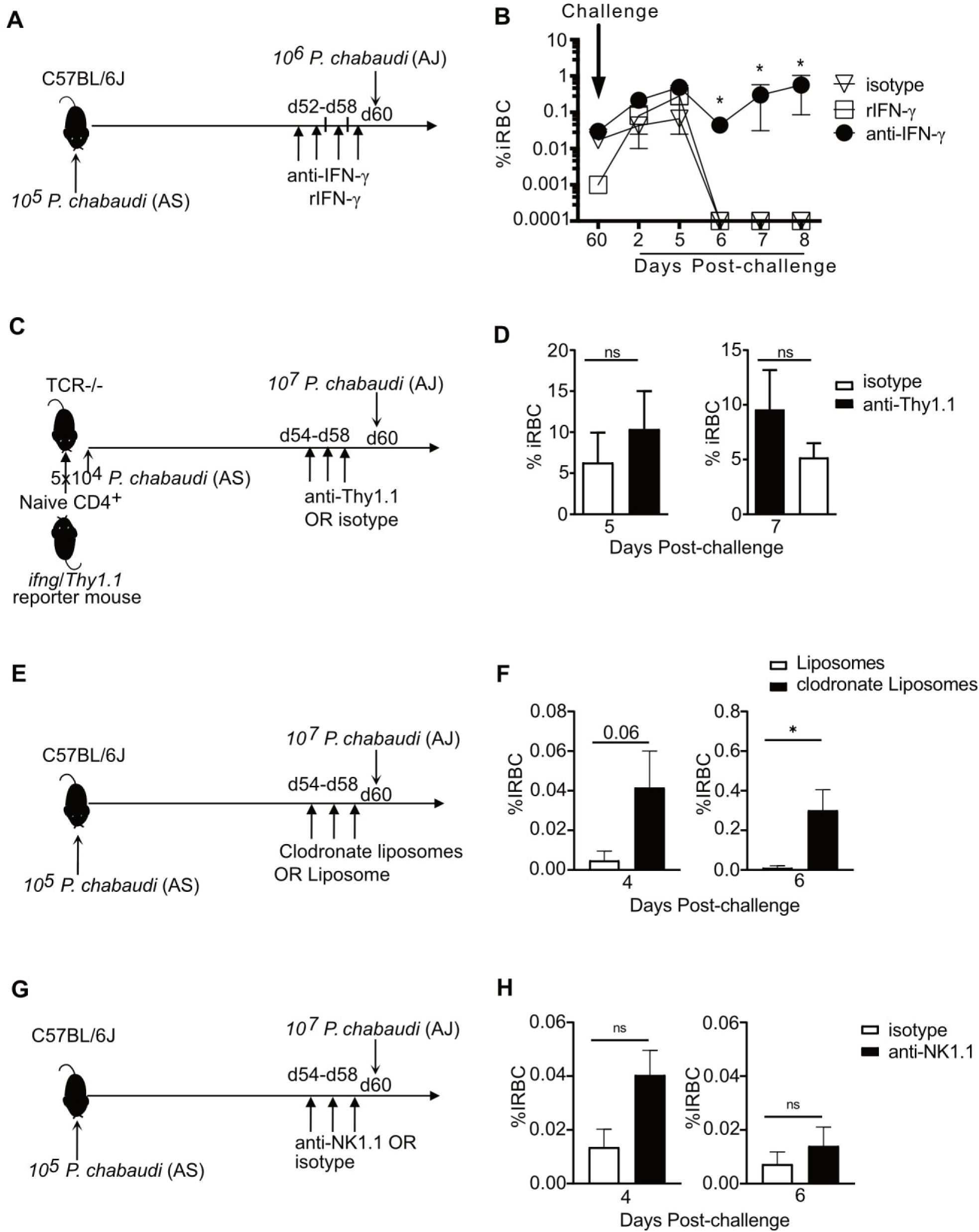
*In vivo* neutralization of IFN-γ prior to re-infection induces loss of premunition, and recombinant IFN-γ restores immunity to secondary infection. (**A**) Schematic showing C57BL/6 mice were infected with *P. chabaudi*. Some animals were administered neutralizing anti-IFN-γ, or recombinant IFN-γ (rIFN-γ), every other day from day 52 to d60 p.i.. Mice were then challenged with heterologous *P. chabaudi* AJ at day 60 post primary infection. (**B**) Graph showing parasitemia measured days post-challenge. (**C**) CD4 T cells from uninfected *Ifng*/*Thy1.1* Knock-In mice were transferred into TCR^-/-^ mice, which were then infected with *P. chabaudi* AS. Some animals were administered anti-Thy1.1, every other day from day 54 to d58 p.i.. Mice were then challenged with heterologous *P. chabaudi* AJ at day 60 post primary infection. (**D**) Graphs showing parasitemia on days 5 and 7 of heterologous challenge of TCR^-/-^ animals. (**E**) C57BL/6 mice were infected with *P. chabaudi*. Some animals were administered with clodronate liposomes or liposome every day from day 55 to d58 p.i.. Mice were then challenged with heterologous *P. chabaudi* AJ at day 60 post primary infection. (**F**) Graph showing parasitemia. (**G**) C57BL/6 mice were infected with *P. chabaudi*. Some animals were administered neutralizing anti-NK1.1 every other day from day 54 to d58 p.i.. Mice were then challenged with heterologous *P. chabaudi* AJ at day 60 post primary infection. (**H**) Graph showing parasitemia. Data are representative of five animals per group. Error bars represent SEM, n.s., not significant, *p < 0.05, Student’s *t* test.

We next tested the hypothesis that *Ifng*^+^ T cells contribute to a state of premution that exists before re-infection. In order to do this, we adoptively transferred naïve CD4^+^ T cells from the *Ifng*/Thy1.1 knock-in reporter mice into TCR^-/-^ recipients. Recipient mice were then infected with *P. chabaudi* AS and treated with anti-Thy1.1 at day 54-58 p.i. to deplete surviving *Ifng/*Thy1.1^+^ cells generated in the infection (**Fig 5C**). *Ifng/*Thy1.1^+^ cells were completely depleted as shown in **Supplemental Fig. 1**; however, other T cells retained the ability to make IFN-γ quickly upon restimulation, as shown by intracellular cytokinestaining. Parasitemia at days 5 and day 7 of the second heterologous infection of TCR^-/-^ did not show a significant difference, but the interpretation is not clear. Either this result means that IFN-γ from non-depleted T cells controls parasitemia from re-infection, or the critical IFN-γ does not come from T cells (**Fig 5D**).

Therefore, we tested the contribution of macrophages and NK cells for a contribution to premunition. C57BL/6J mice were infected with *P. chabaudi* AS followed by heterologous re-infection on day 60 p.i.. As diagrammed, Some animals from each group were administered either clodronate liposomes for macrophage depletion (**Fig 5E**) or anti-NK1.1 antibody for NK cell depletion (**Fig 5G**) before heterologous challenge. The mice treated with clodronate liposomes before re-infection had significantly higher parasitemia compared to control mice treated with liposomes (**Fig 5F**). In contrast, NK cell depletion before re-infection did not significantly affect the parasitemia (**Fig 5H**). These data indicate that both IFN-γ and macrophages are involved in the immune mechanism mediating protection from reinfection during the persistent phase of *P. chabaudi* infection, suggesting Th1-like immunity.

### Persistent Infection decreases memory T cell numbers but maintains *Ifng*+ Teff

In order to test the effect of persistent parasite on maintenance of T cell numbers and *Ifng* accessibility, residual parasitemia was cleared using an anti-malarial drug before measurements. To label *Plasmodium*-specific T cells, *in vivo* BrdU labelling was performed (days 3-10 or 20-30 p.i.) in infected IFN-γ reporter mice, followed by elimination of persistent parasitemia using chloroquine treatment (CQ; days 30-34 p.i., **Fig 6A**). Surviving BrdU^+^ T cells were detected on day 60 p.i. Persistently infected (-CQ) animals had significantly fewer surviving BrdU^+^ CD4^+^ T cells at day 60 p.i., as compared to persistently infected but chloroquine-treated (+CQ) animals (**Fig 6B**). We also tested the effect of the large initial peak of parasite on Tmem survival. However, results were similar whether we tracked T cells that had proliferated during the first peak of T cells (d3-10), or during a period (d20-30) when parasitemia is controlled to below, a still detectable, 1% (7), and less T cell proliferation (**Fig 6C**). Although there were roughly equal fractions and numbers of Teff and Tmem surviving in the –CQ group, the +CQ group showed a significantly larger Tmem (CD127^hi^) than Teff (CD127^-^) population (**Fig 6D**). The increased presence of Tmem at day 60 also occurred in +CQ animals, when the Tmem precursor cells had been generated during days 20-30 p.i. (**Fig 6E**). The fraction of *Ifng^+^* Tmem was more than *Ifng*^+^ cells within Teff (**Fig 6F**). *Ifng* expression per cell (as measured by geometric MFI, far right) was higher in Teff, but not Tmem, in persistently infected animals (-CQ). Together, these data support and extend previous reports that chronic infection reduces survival of CD4 memory T cells (28), and that chronic infection maintains cytokine expression in *P. chabaudi* (18), specifically *Ifng* at the transcriptional level in effector T cells. Therefore, we determined to look at a larger group of Th1 cytokines.

**Figure 6.**
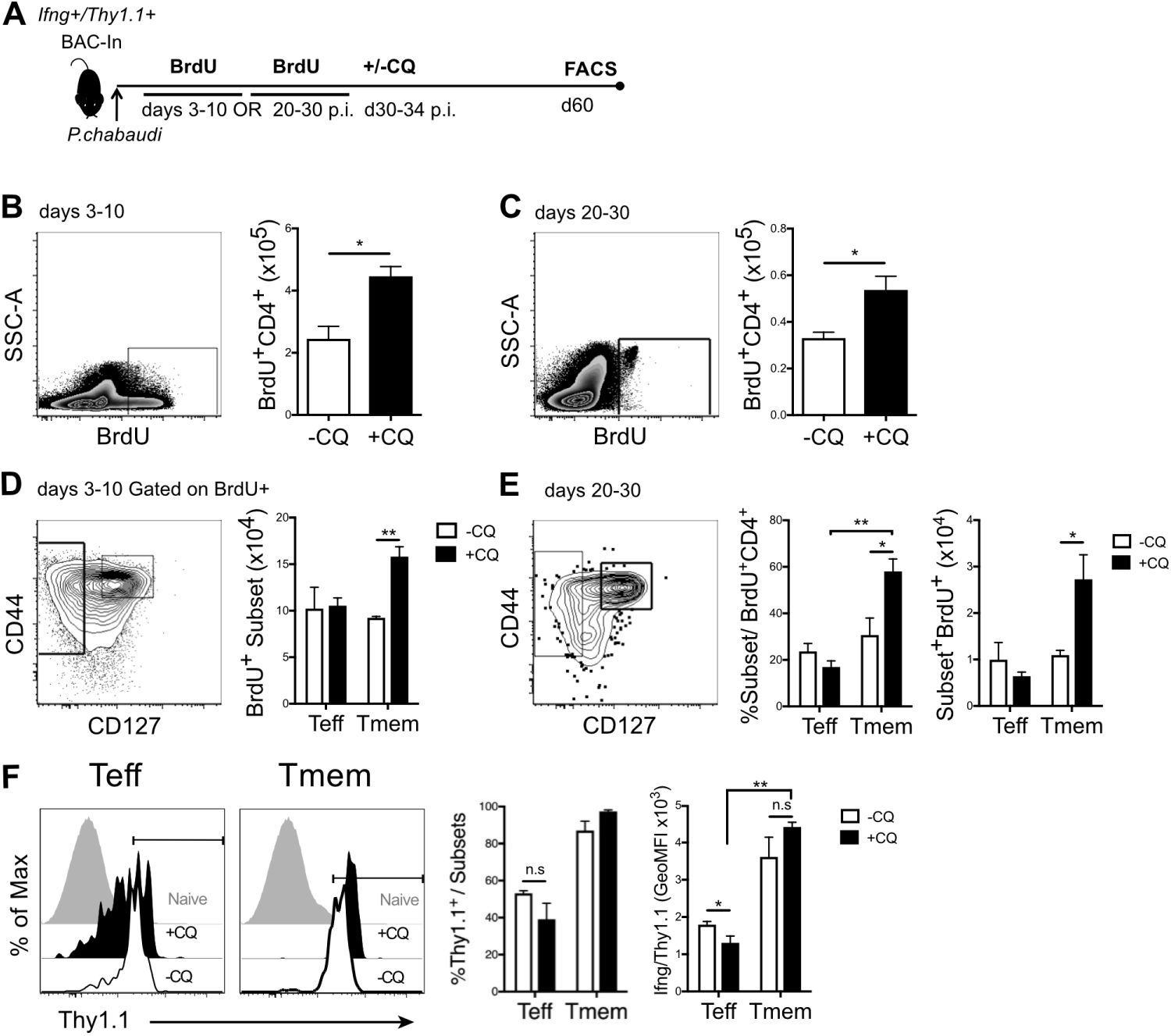
Chronic infection impacts Tmem survival but enhances *Ifng* expression. (**A**) Schematic showing *Ifng*/Thy1.1 BAC-In reporter mice were infected with *P. chabaudi* and given BrdU from days 3-10 or 20-30 p.i.. Some animals were treated with Chloroquine (+CQ, days 30-34 p.i.) to clear residual parasitemia. Splenocytes were harvested at day 60 p.i. for flow cytometry analysis of survival and *Ifng*/Thy1.1 expression. **(B-C)** Zebra plots and graphs showing number of BrdU^+^CD4^+^ T cells in animals receiving **(B)** BrdU days 3-10 or **(C)** 20-30 p.i. in +CQ and -CQ groups. **(D-E)** Contour plots and graphs showing fraction of Teff (CD127**^-^**) and Tmem (CD44^hi^CD127^hi^) out of BrdU^+^ CD4 T cells and number of total BrdU^+^ Teff and Tmem in animals receiving **(D)** BrdU days 3-10 or **(E, F)** 20-30 p.i. in +CQ and -CQ groups. (**F**) Histograms and graph showing the fraction of *Ifng*/Thy1.1^+^ within, and the level (gMFI) of *Ifng*/Thy1.1 expression on, BrdU^+^ Teff and Tmem in +CQ and -CQ control groups. Error bars represent SEM, n.s., not significant. *p < 0.05, **p < 0.01, Student’s *t* test.

### Persistent infection promotes IFN-γ^+^TNF^+^IL-2^-^ T cells

To identify the cytokines produced by effector and memory T cells surviving into the phase of premunition, and their dependence on persistent parasite, we measured IFN-γ, TNF, and IL-2 production by BrdU^+^ T cells in the spleen using intracellular cytokine staining after BrdU labelling from days 20-30 (as in Fig 5C and 5E). Gating on CD127^-^ BrdU^+^ Teff, it was clear that Teff cytokine production upon restimulation was severely limited in persistent infection (**Fig 7A**). Despite in vivo *Ifng* expression, there was only a slight shift in IFN-γ intracellular staining, with clear peaks for TNF and IL-2 only in the drug-treated group. Boolean gating analysis of the distribution of cytokines produced by each Teff population is represented in pie charts and histograms (**Fig 7B**). Teff in persistent infection had a significantly lower fraction of triple and double cytokine producers (IFN-γ^+^TNF^+^IL-2^+^, IFN-γ^-^TNF^+^IL-2^+^) than +CQ Teff. In contrast, BrdU^+^ Tmem from persistent infection exhibited significant IFN-γ and IL-2 intracellular cytokine staining above background (**Fig 7C**). Qualitative differences in cytokine combinations expressed between Teff and Tmem were also noted. IL-2 single producers and IFN-γ^+^IL-2^+^TNF^-^ were detected in Tmem but not in Teff. Single producers of IFN-γ were higher in Tmem than Teff, and these cells were also slightly and significantly increased in –CQ Tmem. The higher IFN-γ protein levels in Tmem (**Fig 7D**) compared to Teff (**Fig 7A**) confirms results seen in the transcriptional readout from the *Ifng* reporter presented in **Fig 6F**.

**Figure 7.**
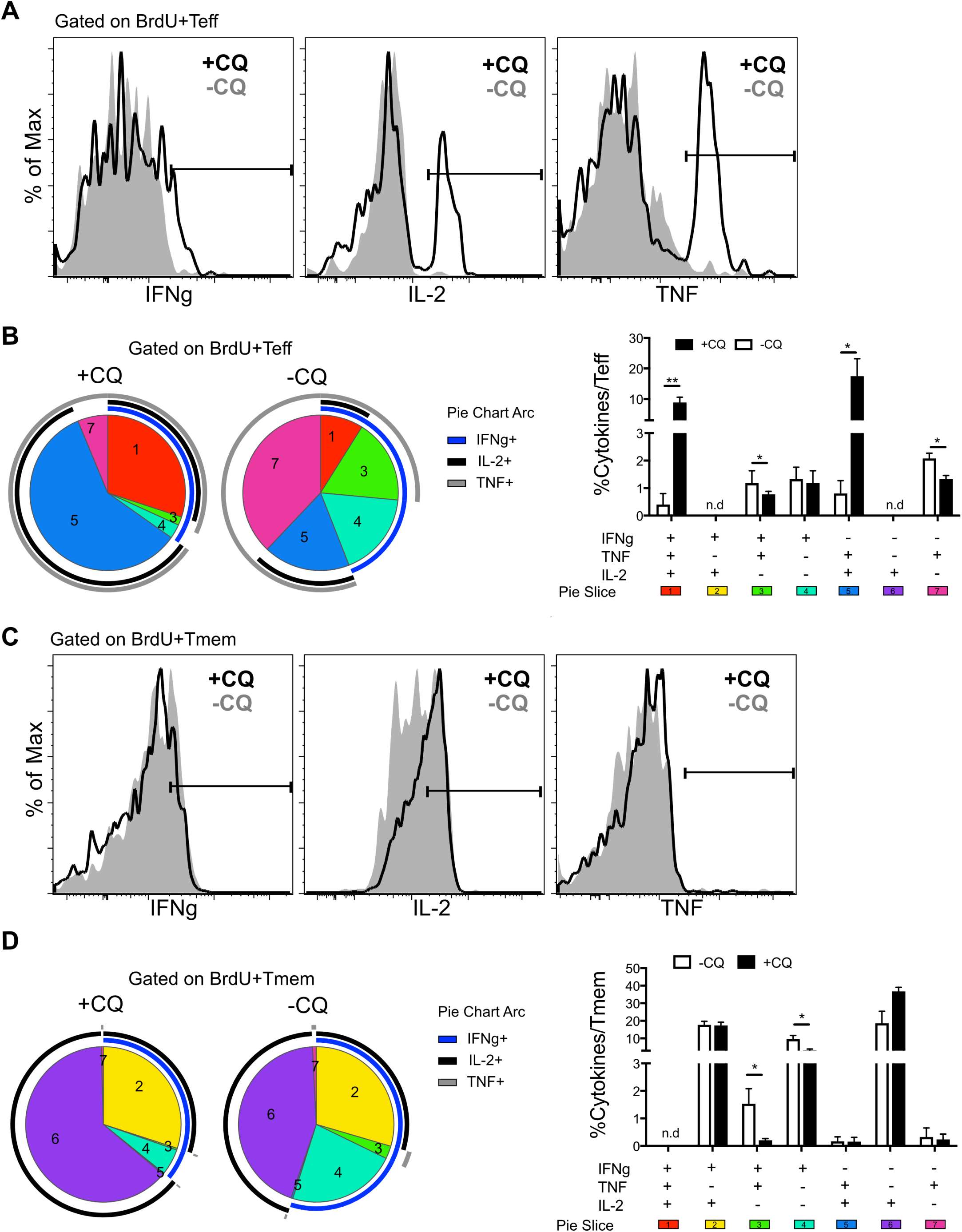
Tmem make highest cytokine levels and persistent infection promotes IFN-γ+TNF+IL-2-T cells, though it reduces Teff multi-cytokine producers. *Ifng*Thy1.1 BAC-In reporter mice were infected with *P. chabaudi* and given BrdU from days 20-30 p.i. Some groups were treated with Chloroquine (+CQ, days 30-34 p.i.) to cure residual parasitemia. Splenocytes were harvested at day 60 p.i. for intracellular cytokine staining. Histograms showing the fraction of IFN-*γ*, IL-2 and TNF production in (**A**) Teff and (**C**) Tmem in +CQ (dark line) and –CQ (gray line). Pie slice representing combination of markers are shown (**B** and **D**). Bar graphs showing Boolean gating analysis of all possible combinations of IFN-*γ*, IL-2, and TNF production within (**B**) Teff and (**D**) Tmem in +CQ (black bar) and -CQ (white bar). Arc lines around the pie represents the fraction of IFN-γ^+^ (blue), IL-2^+^ (black), and TNF^+^ (gray). Data are representative of three independent experiments with four animals per group. Error bars represent SEM, n.d., none detected. p < 0.05, **p < 0.01, Student’s *t* test.

The finding that correlates with the protection maintained by persistent infection, i.e. in – CQ animals compared to treated mice (+CQ), is that IFN-γ^+^TNF^+^IL-2^-^ and IFN-γ single producers were significantly increased among Tmem in persistently infected animals compared to treated. In previous work, a CD62L^lo^ IFN-γ^+^TNF^+^IL-2^-^ population was also detected as increased in MSP-1 specific TCR Tg T cells from chronically-infected-compared to treated-animals at day 60 p.i. (13). This interesting subset appear to be present within both CD127^-^ Teff and Tmem.

## Discussion

In this study, we studied two time periods of declining protection from *Plasmodium* infection. In addition to loss of premunition as parasite disappears after a few months, protection wanes again measurably by day 200 p.i. (29). T cells have been shown to be affected by persistent infection more than Antibody during both periods (13, 15). Our data add to these previous studies and flesh out our understanding of the effect of persistent infection on T cells both as the parasite declines, and then later as the T cell response diminishes enough for parasite to be able to grow again.

Overall, our findings support the interpretation that the promotion of *Ifng^+^* Teff by chronic infection enhances immunity. In the later time period, we show that *Ifng*^+^ T cells in the spleen decay precipitously once the period of premunition is over and then Teff become undetectable, even in the spleen, by day 200, coinciding with the second reduction in protection by day 200 (15). The kinetics of decay of Teff numbers after day 60, in combination with our previous observation that Teff from *P. chabaudi* infection protect better than Tmem also indicate that Teff decline is likely the cause of loss of immunity after day 60. Recombinant IFN-γ administered systemically in the days before re-infection, has been shown to enhance long-term protection in *P. chabaudi* infection, specifically promoting sterile immunity upon re-infection at day 200 (17, 30). Our previous work and that of others shows that there are few if any fully Th1 committed (T-bet^hi^) T cells generated in *P. chabaudi* infection (16, 18, 31). For example, 94% of *Ifng*/Thy1.1^+^ BAC-In Teff transferred from infected donors (day 8 p.i.) to infection-matched recipients lose *Ifng* accessibility (expression of BAC-In *Ifng*/Thy1.1) by day 60 (18). Importantly, the disappearance of Teff detectable in the blood by day 200, and the *Ifng*^+^ Teff by day 100 p.i., corresponds to the timeframe when loss of protection, or re-appearance of susceptibility to *P. chabaudi* parasitemia upon re-infection, was documented by do Rosario *et al.* as between day 120 and day 200 p.i. (15). Together, these three studies suggest that decay of *Ifng*+ Teff explains the second decay of protection, but this does not fully explain the enhanced protection seen during persistent infection in the earlier phase.

In addition to improving our understanding of the long-term decay of immunity, this study identifies some changes in T cell function specifically downstream of low-level persistence of parasite. The literature suggests that while maintenance of antigen-specific T cells after stimulation does not require MHCII, survival and protective CD4 T cell functions can be improved by persistent antigen (32–34). In addition, while resting Tcm are capable of providing some long-term immunity against *Leishmania major*, a prototypical persistent pathogen, persistence of the pathogen, actually promotes solid immunity to reinfection (35). However features of persistent stimulation that contribute to protection have been elusive. Multifunctional Th1 cells (IFN-γ^+^TNF^+^IL-2^+^) have been shown to correlate with protection after vaccination to *Leishmania* (36), though it has not been established if these are long-lived Tmem or short-lived Teff. Our work suggests that in cured persistent infection, these cells are largely Teff. As chronic infection enhances protection to both *Leishmania* and *Plasmodium* (7, 37), T cell multi-functionality could have been the correlate of immunity during premunition (14). However, persistently stimulated (-CQ) Teff have a significantly lower fraction of triple cytokine producers (IFN-γ^+^TNF^+^IL-2^+^) compared to rested T cells (+CQ). On the other hand, an IFN-γ^+^TNF^+^IL-2^-^ CD127^+/-^ T cell subset has been shown, using an adoptive transfer system, to be increased in persistent *P. chabaudi* infection compared to treated animals (13). Here, we show that in polyclonal response these IFN-γ^+^ T cells are also are promoted by persistence of parasite, and that these cells contain both Teff and Tmem. Both intracellular cytokine staining and reporter data show that more Tmem than Teff produce cytokines; however, triple cytokine producers on day 60 are confined to BrdU^+^CD127^-^ Teff. This surprising result suggests that triple producing T cells, reported to correspond to protection in several models, are short-lived effector T cells. CD127 expression and longevity will have to be tested in other models to confirm that this is true in other infections. However, this population is also curtailed by persistent infection, at least when measured by intracellular cytokine staining, despite persistent infection being protective, suggesting multi-functional cytokine producing CD4 T cells are not a great correlate of protection in the case of premunition.

In order to determine the role of IFN-γ on the first phase of protection, where *Ifng*^+^ Teff are promoted by persistent infection, we tested the effect of neutralizing IFN-γ on the premunition phase. Neutralization of IFN-γ before infection eliminated the improved protection provided by persistent infection, or premunition. In addition, we tested the contribution of NK cells and macrophages to protection in this phase. Depletion of phagocytic cells, but not NK cells also reduced the effect of premunition and prolonged re-infection challenge parasitemia. Each of these cell types may contribute to IFN-γ production. NK cells and macrophages can contribute for example in *Salmonella* infection (38). Neutrophils and microglia have measurable effects on *Toxoplasma* infection (39) *in vivo*. While training of phagocytes is reported to occur by persistent viral infection (40), it is challenging to determine from these experiments if Th1 cytokines associated with persistent infection are primarily critical in the period before re-infection for training, or also during re-infection. Each of the experiments one could envision for this test has caveats, which is why we performed several. Both antibodies and clodronate have the potential to persist for weeks, and it would be difficult to establish the threshold at which they are no longer potent. Therefore the limited conclusion is that Th1 type responses, and not just antibody, are critical for rapid clearance of repeat infection during persistent infection. This assertion is also supported by our recent demonstration that STAT3 deficiency in T cells shifts the response towards Th1 and protects in *Plasmodium* re-infection, (16). It will be important to use technology to attempt to discriminate between the effects of innate cell conditioning during premunition and adaptive cell memory reactivation upon re-infection.

Here we also showed in a physiological setting that Tmem survive and contain a population of Tem that maintain Ifng expression. We and others have shown that this CD27^-^ population is propagated by the other populations (13, 41). This confirms and extends our previous work using *P. chabaudi* MSP1-specific B5 TCR Tg that showed all Tmem subsets, including Tem, survive longer than Teff (20). However, there is definitely an effect of persistent infection on Tmem number. This could be interpreted as a deficiency in Tmem in chronic infection, except that immunity to reinfection is actually better in low-level persistent infection than after cure of the same infection (5). Similar experiments have been done using naïve adoptively transferred B5 TCR Tg Teff and showed a similar decrease in Tmem in persistently infected mice compared to treated (13). Our previous work on mechanisms of Tem survival shows that in addition to maintenance of Tem numbers in the absence of antigen, adoptively transferred Tem can proliferate in response to the low-level persistent infection late in *P. chabaudi* infection, though only the poorly protective CD27^+^ Tem^Early^ subset expand in number as a result (14). Therefore, the present data suggest that persistent infection promotes Ifng persistence in short-lived *Ifng*^+^ Teff but not Tem and that this constitutes an important facet of premunition.

## Materials and Methods

### Mice and Infections

*Ifng*/Thy1.1 BAC-In or *Ifng*/Thy1.1 knock-In mice were a kind gift of Casey Weaver (University of Alabama, Birmingham, AL). C57BL/6 and TCRβ/δ^-/-^ (B6.129P2-Tcrb^tm1Mom^Tcrd^tm1Mom^) mice were purchased from the Jackson Laboratory (Bar Harbor, ME). The BAC-In *Ifng*/Thy1.1 construct is a faithful marker of the accessibility of the *Ifng* locus, representing a fraction of the total cells that stain positive for IFN-γ protein using intracellular cytokine staining (22). The construct includes an SV40 polyA tail to stabilize the reporter at the RNA level, so expression has been shown to last at least 40 days after its first expression (22). These mice were maintained in our specific pathogen-free animal facility with *ad libitum* access to food and water. Mice 6–12 weeks old were infected i.p. with 10^5^ *Plasmodium chabaudi chabaudi* AS (courtesy of Jean Langhorne, Francis Crick Institute, London, UK). For protection assays, C57BL/6 mice were infected i.p. with 10^5^ *P. chabaudi* AS and challenged with 10^6^ *P. chabaudi* AJ (MR4, Manassas, VA) infected erythrocytes. *P. chabaudi* was maintained as described elsewhere (7, 42). Parasites were counted by light microscopy in thin blood smears stained with Giemsa (Sigma-Aldrich, St. Louis, MO). All animal studies were carried out in accordance with the protocol as approved by the University of Texas Medical Branch Institutional Animal Care and Use Committee.

### In vivo assays

For *in vivo* labeling with 5-bromo-2’-deoxyuridine (BrdU; Sigma), BrdU was given in drinking water (0.8mg/ml) on days 3-10 or 20-30 p.i. In some experiments, mice were treated with a dose of 50 mg/kg of the antimalarial drug chloroquine (CQ, i.p.) on days 30–34 post-infection (p.i.). C57BL/6 mice were infected with *P. chabaudi* and then treated with four doses of (0.5mg/mouse) of depleting mAbs against IFN-γ or isotype control antibody (H22; Thermo Fisher Scientific, Waltham, MA) or (15 ng/mouse) of recombinant-mouse IFN-γ (rIFN-γ expressed in *E. coli*; PeproTech, Rocky Hill, NJ), i.p. every 2 d starting 52 days p.i. as previously established (17). For NK cells depletion, mice were injected with either an anti-NK1.1-depleting antibody or isotype control antibody (PK136, BioXCell Lebanon, NH). Clodronate liposomes have been used for macrophages depletion in several immunology studies to investigate on their functions (43, 44). For macrophages depletion, 100 µL Clodronate liposomes (0.5mg) or liposomes were administrated with 5 doses every day starting 55 days to chronic infected mice before reinfection.

In some experiments, Ifng/Thy1.1 CD4 T cells from uninfected 5 week old animals were MACS sort purified and transferred into TCRβ/δ-/- (TCR^-/-^) recipients, which were infected and then treated three times with anti Thy1.1 (250ug/mouse i.p., Clone 19E12, BioXCell, West Lebanon, NH) every other day between days 56-60 p.i., similar to the experiment described in (45).

### Flow cytometry

Blood was collected by tail bleeding with a heparin-coated pipette tip and incubated in RBC Fix/Lysis buffer (eBioscience, San Diego, CA) for 10 min at room temperature before washing with PBS containing 2% FBS and 0.1% sodium azide (FACS buffer; Sigma Aldrich, St. Louis, MO). Cells were then stained with the following antibodies: anti-CD90.1 (30-H12), anti-CD44 (IM7), and anti-CD27 (LG-7F9), and the following combinations of markers: PerCP/efluor710, allophycocyanin (APC)/efluor780 and APC-conjugated antibodies (all from eBioscience), anti-CD127 (A7R34) PE-Cyanine-5, anti-CD62L (MEL-14) Brilliant Violet 605, and CD4 (RM4-5)-Brilliant Violet 650 (BioLegend, San Diego, CA).

Single-cell suspensions from spleens were prepared in HEPES-buffered HBSS (Life Technologies, Thermo Fisher Scientific, Waltham, MA) and prepared with RBC lysis buffer (eBioscience) before washing with FACS buffer. Cells were then stained with Fc receptor blocking antibody (Clone 2.4G2, BioXCell), followed by surface staining as described above in blood. Cells were stained using the FITC BrdU flow kit (B44, BDbiosciences) and analyzed by flow cytometry according to the manufacturer’s protocol.

For intracellular staining, splenocytes (5 x 10^6^ per ml) were stimulated with PMA (50 ng/mL) and ionomycin (500 ng/mL) at 37°C for 5 hours in complete Iscove’s Media (cIMDM). Brefeldin A (10 μg/mL) was added for the last two hours (all from Sigma). Cells were processed for surface staining as described above and fixed with 2% paraformaldehyde (Sigma-Aldrich) followed by permeabilization using 1X Permeabilization buffer (BDbiosciences). Cells were washed 3 times in permeabilization buffer and incubated for 40 minutes with anti-IFN-γ-FITC (XMG1.2), IL-2-CF594 (JES6-5H4), and TNF-PE/Cy7 (MP6-XT22) or isotype controls (all from eBioscience). Cells were collected on a LSRII Fortessa using FACSDiva software (BDbiosciences) and analyzed in FlowJo (version 9.7, Tree Star, Ashland, OR). Compensation was performed in FlowJo using single-stained splenocytes (with CD4 in all colors). Boolean gating analysis was performed, and the distribution of cytokines was analyzed with Spice version 6 (46) and Prism, GraphPad (version 7, La Jolla, CA). Pie charts show fractions of cytokine producers only. Data from each mouse was analyzed, and averages and SEM were calculated for graphs and contour plots. Data from five mice were concatenated to achieve sufficient cell numbers for presentation.

### Statistical analysis

All data are presented as mean +/- SEM. Two-tailed unpaired Student’s *t* test was used (Prism, GraphPad, La Jolla, CA). *p < 0.05, **p < 0.01, ***p < 0.001, n.s. not significant; n.d., none detected. The limit of detection (L.O.D) was calculated using all available data from animals from all experiments that did not receive BrdU, and is defined as the mean of this background x 1.645.

## Acknowledgements

We are grateful to Dr. Jean Langhorne’s curiosity about how long *Plasmodium*-specific memory T cells last for the inspiration to do this work. We would like to thank Laurie Harrington for advice. We acknowledge the support of the UTMB Microbiology and Immunology Flow Cytometry Core and the Animal Research Core.

---

**Supplemental Figure 1.**
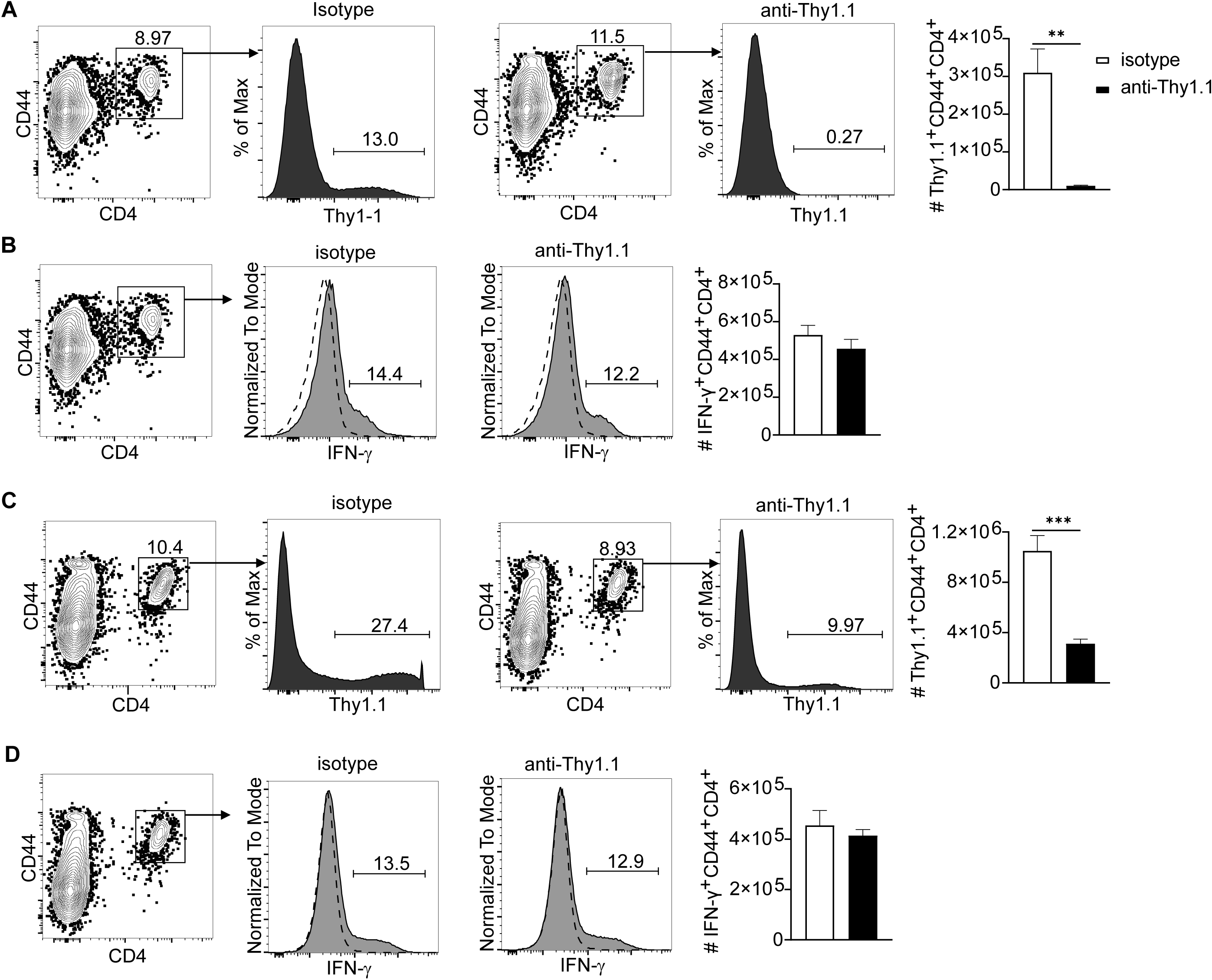
Depletion of *Ifng*-reporter cells and IFN-γ production after stimulation. CD4 T cells from uninfected *Ifng*/Thy1.1 Knock-In mice were transferred into TCR^-/-^ mice, which were then infected with *P. chabaudi* AS. Some animals were administered anti-Thy1.1, every other day from day 54 to d58 p.i.. Splenocytes were harvested at day 60 p.i. or day 20 post challenge for flow cytometry analysis to check the depletion *Ifng*/Thy1.1 Knock-In and for intracellular cytokine staining. Plots, Histograms and graph showing gating and the fraction of (**A**) *Ifng*/Thy1.1 Knock-In cells in CD4^+^CD44 ^hi^ after depletion at day 60 p.i.; (**B**) CD4^+^CD44^hi^ gated IFN-γ intracellular cytokine staining at day 60 p.i.; (**C**) *Ifng*/Thy1.1 Knock-In cells in CD4^+^CD44 ^hi^ after challenge at day 20 p.c.; (**D**) CD4^+^CD44^hi^ gated IFN-γ intracellular cytokine staining after challenge at day 20 p.c., dashed line represents isotype control for intracellular staining. Data are representative of three animals per group. Error bars represent SEM. **p < 0.01, ***p < 0.001 Student’s *t* test.

## Notes

Funding: This work was supported by the NIH National Institute of Allergy and Infectious Diseases Grants R01AI08995304, R01AI135061, R01AI08995304 (RS, VHC, SAI, MMO, KG) R01AI08995304S01 (RS, VHC), F31AI126809 (VHC), James W. McLaughlin Postdoctoral and Predoctoral Fellowships at UTMB (KG, VHC), and the Jeane B. Kempner Fellowships (V.H.C.), and the American Association of Immunologists Careers in Immunology Fellowship (SAI).

